# A protocol for adding knowledge to Wikidata, a case report

**DOI:** 10.1101/2020.04.05.026336

**Authors:** Andra Waagmeester, Egon L. Willighagen, Andrew I Su, Martina Kutmon, Jose Emilio Labra Gayo, Daniel Fernández-Álvarez, Quentin Groom, Peter J. Schaap, Lisa M. Verhagen, Jasper J. Koehorst

## Abstract

Pandemics, even more than other medical problems, require swift integration of knowledge. When caused by a new virus, understanding the underlying biology may help finding solutions. In a setting where there are a large number of loosely related projects and initiatives, we need common ground, also known as a “commons”. Wikidata, a public knowledge graph aligned with Wikipedia, is such a commons and uses unique identifiers to link knowledge in other knowledge bases However, Wikidata may not always have the right schema for the urgent questions. In this paper, we address this problem by showing how a data schema required for the integration can be modelled with entity schemas represented by Shape Expressions. As a telling example, we describe the process of aligning resources on the genomes and proteomes of the SARS-CoV-2 virus and related viruses as well as how Shape Expressions can be defined for Wikidata to model the knowledge, helping others studying the SARS-CoV-2 pandemic. How this model can be used to make data between various resources interoperable, is demonstrated by integrating data from NCBI Taxonomy, NCBI Genes, UniProt, and WikiPathways. Based on that model, a set of automated applications or bots were written for regular updates of these sources in Wikidata and added to a platform for automatically running these updates. Although this workflow is developed and applied in the context of the COVID-19 pandemic, to demonstrate its broader applicability it was also applied to other human coronaviruses (MERS, SARS, Human Coronavirus NL63, Human coronavirus 229E, Human coronavirus HKU1, Human coronavirus OC4).

## Introduction

The COVID-19 pandemic, caused by the SARS-CoV-2 virus, is leading to a burst of swiftly released scientific publications on the matter (1). In response to the pandemic, many research groups have started projects to understand the SARS-CoV-2 virus life cycle and to find solutions. Examples of the numerous projects include outbreak.info (2), VODAN around FAIR data (3), CORD-19-on-FHIR (4) and the COVID-19 Disease Map (5). Many research papers and preprints get published every week and many call for more Open Science (6). The Dutch universities went a step further and want to make any previously published research openly available, in whatever way related to COVID-19 research (7).

However, this swift release of research findings comes with an increased number of incorrect interpretations (8) which can be problematic when new research articles are picked up by main-stream media (9). Rapid evaluation of these new research findings and integration with existing resources requires frictionless access to the underlying research data upon which the findings are based. This requires interoperable data and sophisticated integration of these resources. Part of this integration is reconciliation, which is the process where matching concepts in Wikidata are sought. Is a particular gene or protein already described in Wikidata? Using a shared interoperability layer, like Wikidata, different resources can be more easily linked.

The Gene Wiki project has been linking the different research silos on genetics, biological processes, related diseases and associated drugs (10), creating a brokerage system between the research silos. The project recognises Wikidata as a sustainable infrastructure for scientific knowledge in the life sciences.

In contrast to legacy databases, where data models follow a relational data schema of connected tables, Wikidata (https://wikidata.org/) uses statements to store facts (see Figure 1) (10–13). This model of statements aligns well with the RDF triple model of the semantic web and the content of Wikidata is also serialized as Resource Description Framework (RDF) triples (14,15), acting as a stepping stone for data resources to the semantic web. Through its SPARQL endpoint (https://query.wikidata.org), knowledge captured in Wikidata can be integrated with other nodes in the semantic web, using mappings between these resources or through federated SPARQL queries (16).

**Figure 1:**
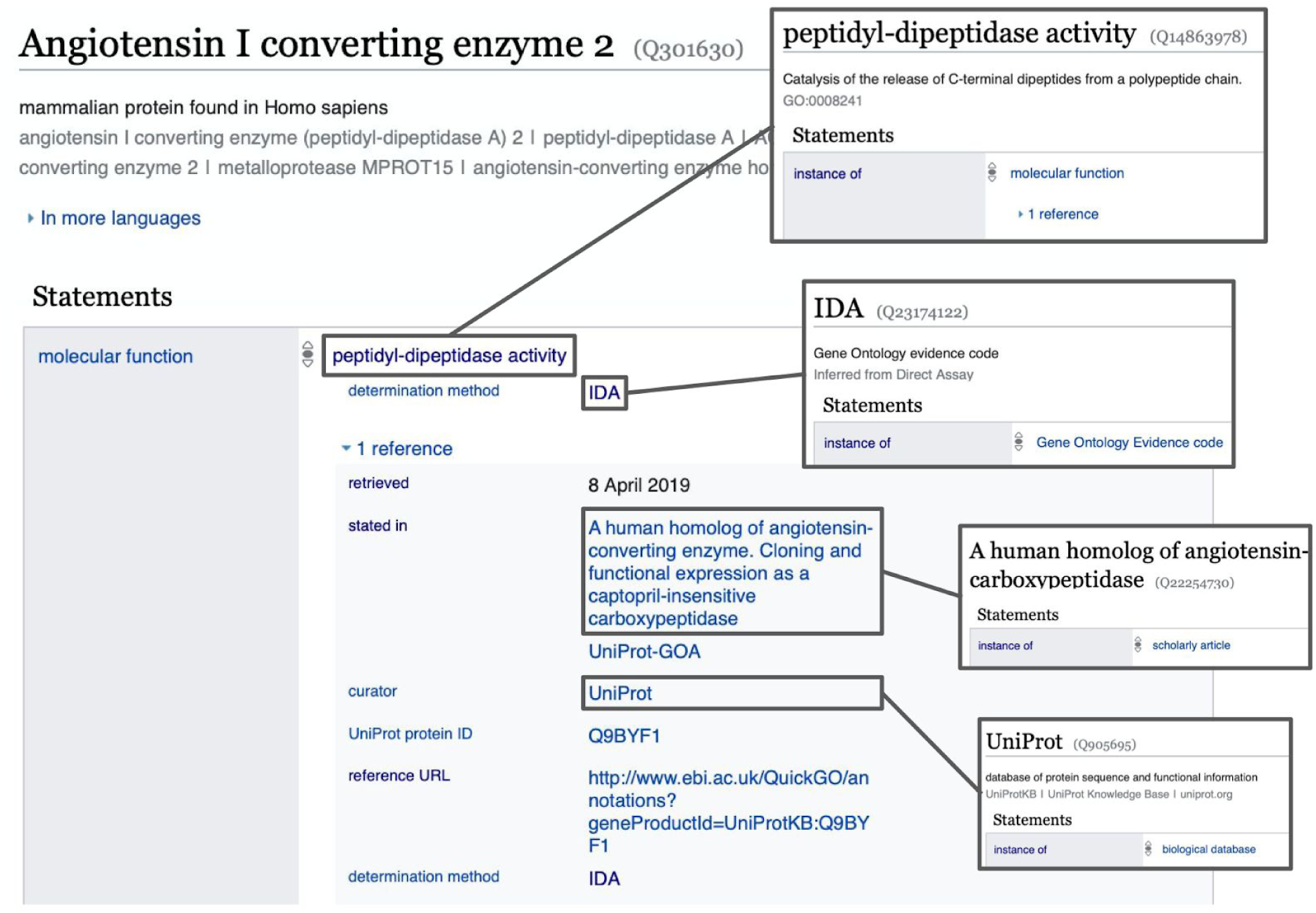
Structure of a Wikidata item, containing a set of statements which are key-value pairs, with qualifiers and references. Here the item for the ACE2 protein is given containing a statement about its molecular function. This molecular function (peptidyl-dipeptidase activity) contains a reference stating when and where this information was obtained.

The Gene Wiki project aligns novel primary data sources with Wikidata in a cycle of consecutive steps where the data schema of the primary source is aligned with the available Wikidata properties. Once the schema is in place, bots are developed to add and regularly update Wikidata with knowledge from the primary resource under scrutiny.

Automated editing of Wikidata simplifies the process, however, the quality control must be monitored carefully. This requires a clear data schema that allows the various resources to be linked together with their provenance. This schema describes the key concepts required for the integrations of the resources we are interested in the NCBI Taxonomy (17), NCBI Gene (18), UniProt (19), the Protein Data Bank (PDB) (20), WikiPathways (21), and PubMed (22). Therefore, the key elements for which we need a model include viruses, virus strains, virus genes, and virus proteins. The first two provide the link to taxonomies, the models for genes and proteins link to UniProt, PDB, and WikiPathways. These key concepts are also required to annotate research output such as journal articles and datasets related to these topics. Wikidata calls such keywords ‘main subjects’. The introduction of this model and the actual SARS-CoV-2 genes and proteins in Wikidata enables the integration of these resources.

This paper is a case report of a workflow/protocol for data integration and publication. The first step in this approach is to develop the data schema. Within Wikidata, Shape Expressions (ShEx) are used as the structural schema language to describe and capture schemas of concepts (23,24). With ShEx we describe the RDF structure by which Wikidata content is made available. These Shapes have the advantage that they are easily exchanged and describe linked data models as a single knowledge graph. Since the Shapes describe the model, they enable discussion, revealing inconsistencies between resources and allow for consistency checks of the content added by automated procedures.

With the model defined, the focus can turn to the process of adding knowledge to Wikidata. In this phase, the seven human coronaviruses (HCoVs), MERS, SARS, SARS-CoV-2 (causing COVID-19), Human Coronavirus NL63, Human coronavirus 229E, Human coronavirus HKU1, and Human coronavirus OC4 (25), can be added to Wikidata. This protocol is finalized by describing how the resulting data schema and data can be applied to support other projects, particularly the WikiPathways COVID Portal.

The Semantic Web was proposed as a vision of the Web, in which information is given well-defined meaning and better enabling computers and people to work in cooperation (26). In order to achieve that goal, several technologies have appeared, like RDF for describing resources (15), SPARQL to query RDF data (27) and the Web Ontology Language (OWL) to represent ontologies (28).

Linked data was later proposed as a set of best practices to share and reuse data on the web (29). The linked data principles can be summarized in four rules that promote the use of uniform resource identifiers (URIs) to name things, which can be looked up to retrieve useful information for humans and for machines using RDF, as well as having links to related resources. These principles have been adopted by several projects, enabling a web of reusable data, known as the linked data cloud (https://lod-cloud.net/), which has also been applied to life science (30).

One prominent project is Wikidata, which has become one of the largest collections of open data on the web (16). Wikidata follows the linked data principles offering both HTML and RDF views of every item with their corresponding links to related items, and a SPARQL endpoint called the Wikidata Query Service. Wikidata’s RDF model offers a reification mechanism which enables the representation of information about statements like qualifiers and references (see also https://www.wikidata.org/wiki/Help:Statements). For each statement in Wikidata, there is a direct property in the `wdt` namespace that indicates the direct value. In addition, the Wikidata data model adds other statements for reification purposes that allow enrichment of the declarations with references and qualifiers (for a topical treatise, see Ref. (31)). As an example, item <u>Q14875321</u>, which represents ACE2 (protein-coding gene in the *Homo sapiens* species) has a statement specifying that it has a chromosome (<u>P1057</u>) with value chromosome X (<u>Q29867336</u>). In RDF Turtle, this can be declared as:

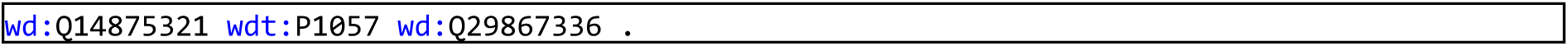

That statement can be reified to add qualifiers and references. For example, a qualifier can state that the genomic assembly (<u>P659</u>) is GRCh38 (<u>Q20966585</u>) with a reference declaring that it was stated (<u>P248</u>) in Ensembl Release 99 (<u>Q83867711</u>). In Turtle, those declarations are represented as (see also Figure 2):

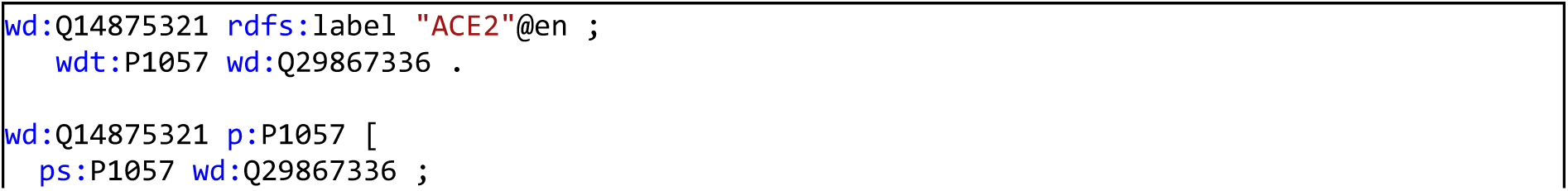

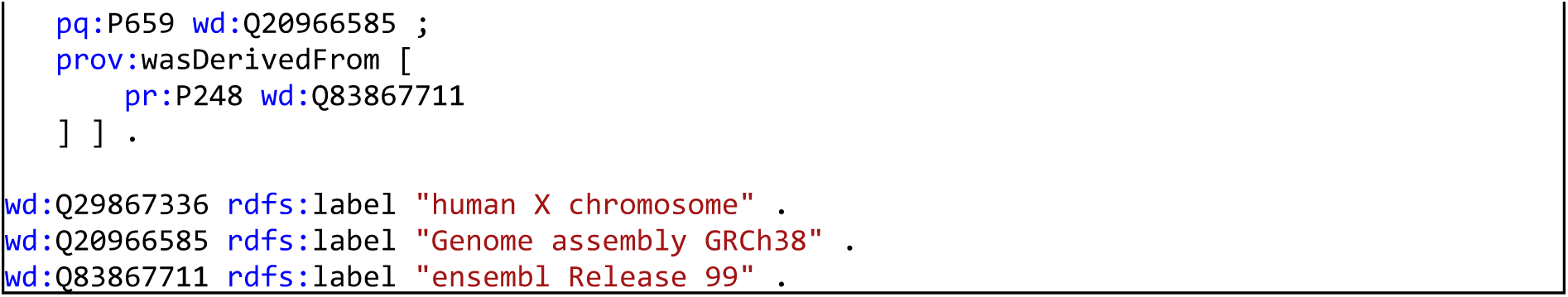

**Figure 2:**
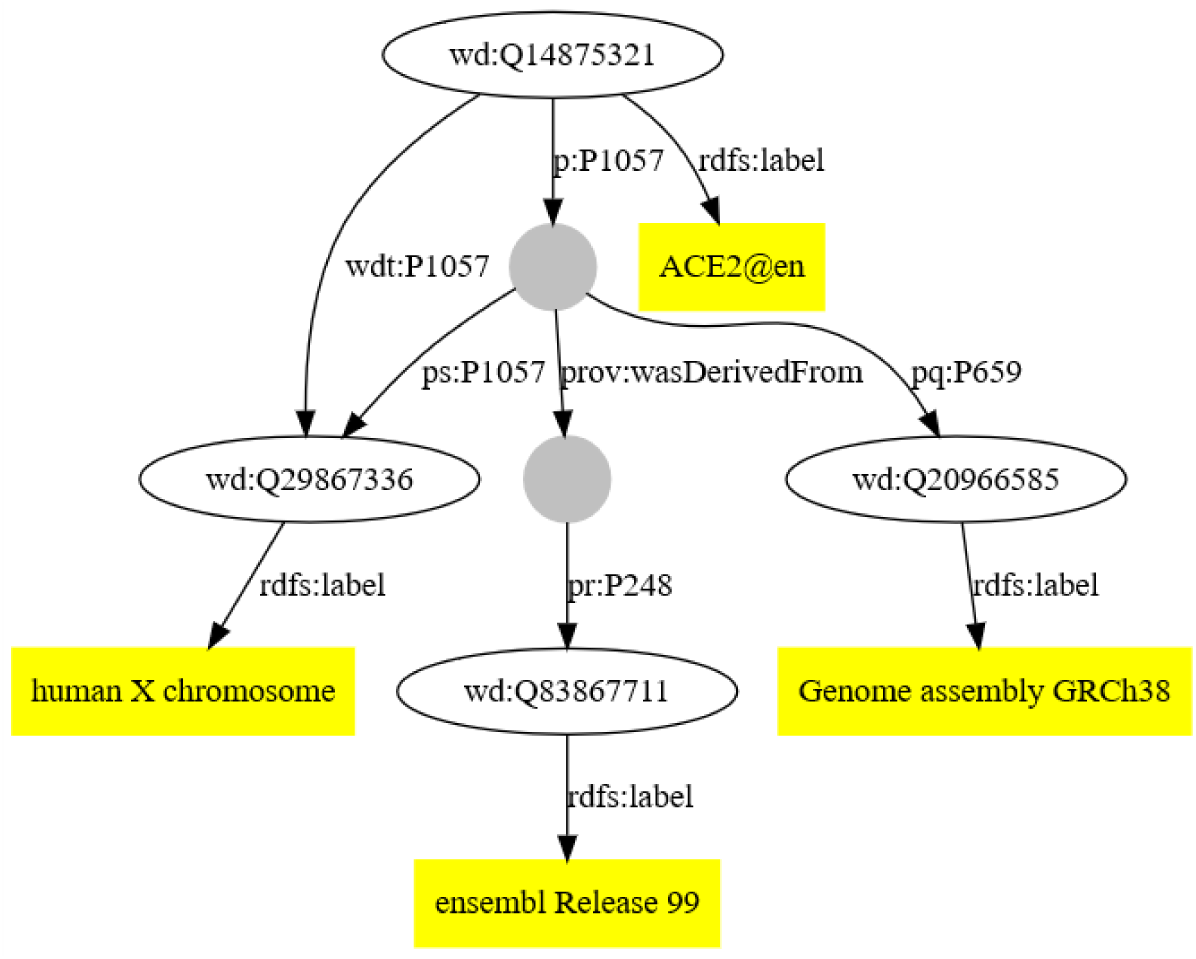
Example of an RDF data model representing ACE2, created with RDFShape (32).

## Methods

### Specifying data models with ShEx

Although the RDF data model is flexible, specifying an agreed structure for the data allows domain experts to identify the properties and structure of their data facilitating the integration between heterogeneous data sources. Shape Expressions were used to provide a suitable level of abstraction. YaShE, http://www.weso.es/YASHE/, a ShEx editor implemented in JavaScript, was applied to author these Shapes (33). This application provides the means to associate labels in the natural language of Wikidata to the corresponding identifiers. The initial entity schema was defined with YaShE as a proof of concept for virus genes and proteins. In parallel, statements already available in Wikidata were used to automatically generate an initial shape for virus strains with sheXer (34). The statements for virus strains were retrieved with SPARQL from the Wikidata Query Service (WDQS). The generated Shape was then further improved through manual curation. The syntax of the Shape Expressions was continuously validated through YaShE and the Wikidata Entity Schema namespace was used to share and collaboratively update the schema with new properties. Figure 3 gives a visual outline of these steps.

**Figure 3:**
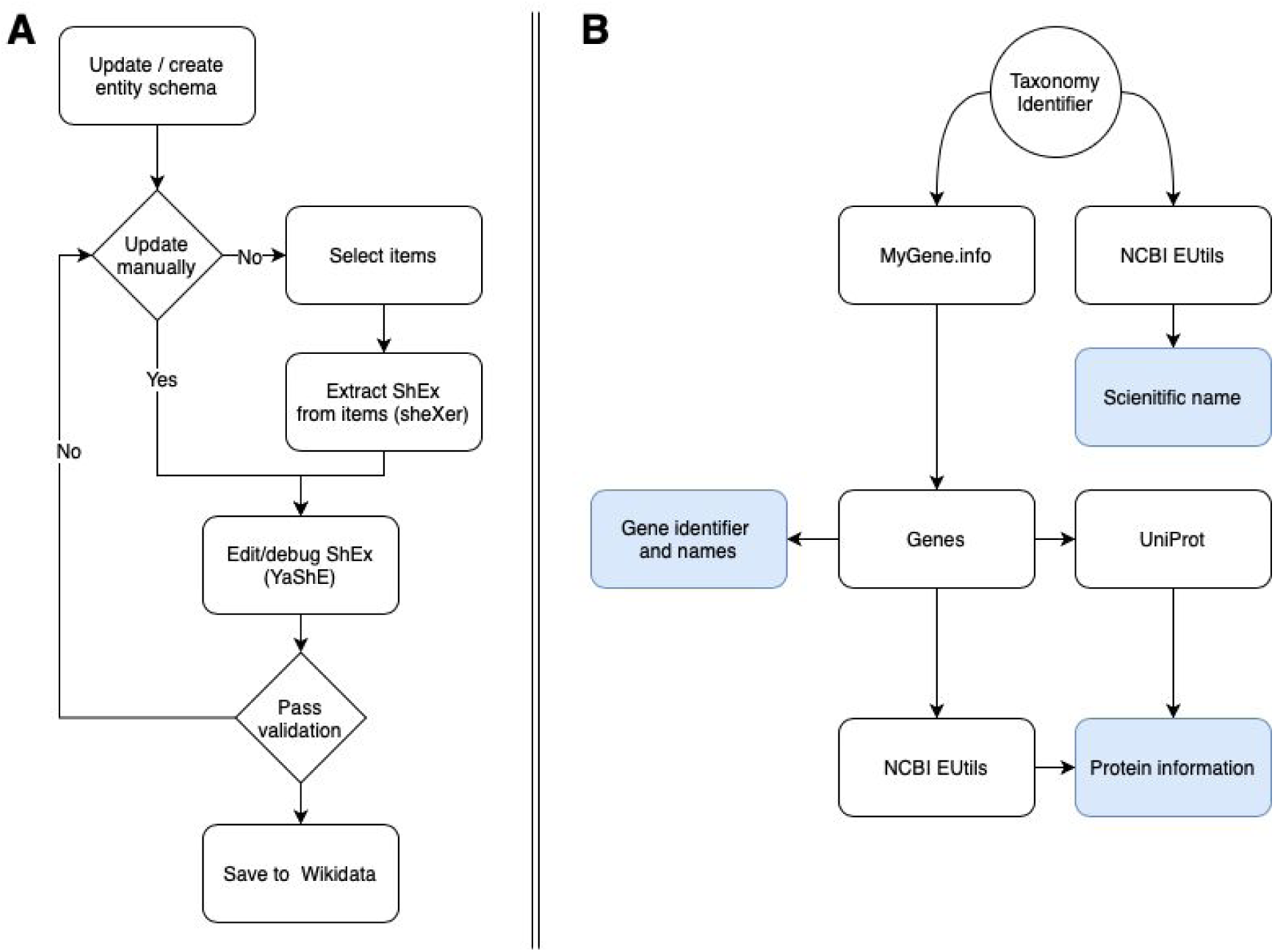
Flow diagram for entity schema development and the executable workflow for the virus gene protein bot. A) The workflow of creating shape expressions. B) The computational workflow of how information was used from various public resources to populate Wikidata.

### Populating Wikidata with Human Coronavirus Data

The second step in our workflow is to add entries for all virus strains, genes and their gene products to Wikidata. This information is spread over different resources. Here, annotations were obtained from NCBI EUtils (35), Mygene.info (36), and UniProt, as outlined below. Input to the workflow is the NCBI Taxonomy identifier of a virus under scrutiny. (*e*.*g*. <u>2697049</u> for SARS-CoV-2). The taxon annotations are extracted from NCBI EUtils. The gene and gene product annotations are extracted from mygene.info and the protein annotations are extracted from UniProt using the SPARQL endpoint at (https://sparql.uniprot.org/).

Genomic information from seven human coronaviruses (HCoVs) was collected, including the NCBI Taxonomy identifiers. For six virus strains, a reference genome was available and was used to populate Wikidata. For SARS-CoV-1, the NCBI Taxonomy identifier referred to various strains, though no reference strain was available. To overcome this issue, the species taxon for SARS-related coronaviruses (SARSr-CoV) was used instead, following the practices of NCBI Genes and UniProt.

### NCBI Eutils

The Entrez Programming Utilities (EUtils) is the application programming interface (API) to the Entrez query and database system at the National Center for Biotechnology Information (NCBI). From this set of services the scientific name of the virus under scrutiny was extracted (e.g. “Severe acute respiratory syndrome coronavirus 2”).

### Mygene.info

Mygene.info is a web service which provides a REST API that can be used to obtain up-to-data gene annotations. The first step in the process is to get a list of applicable genes for a given virus by providing the NCBI taxon id. The following step is to obtain gene annotations for the individual genes from mygene.info through http://mygene.info/v3/gene/43740571. This results in the name and a set of applicable identifiers (Figure 4).

**Figure 4:**
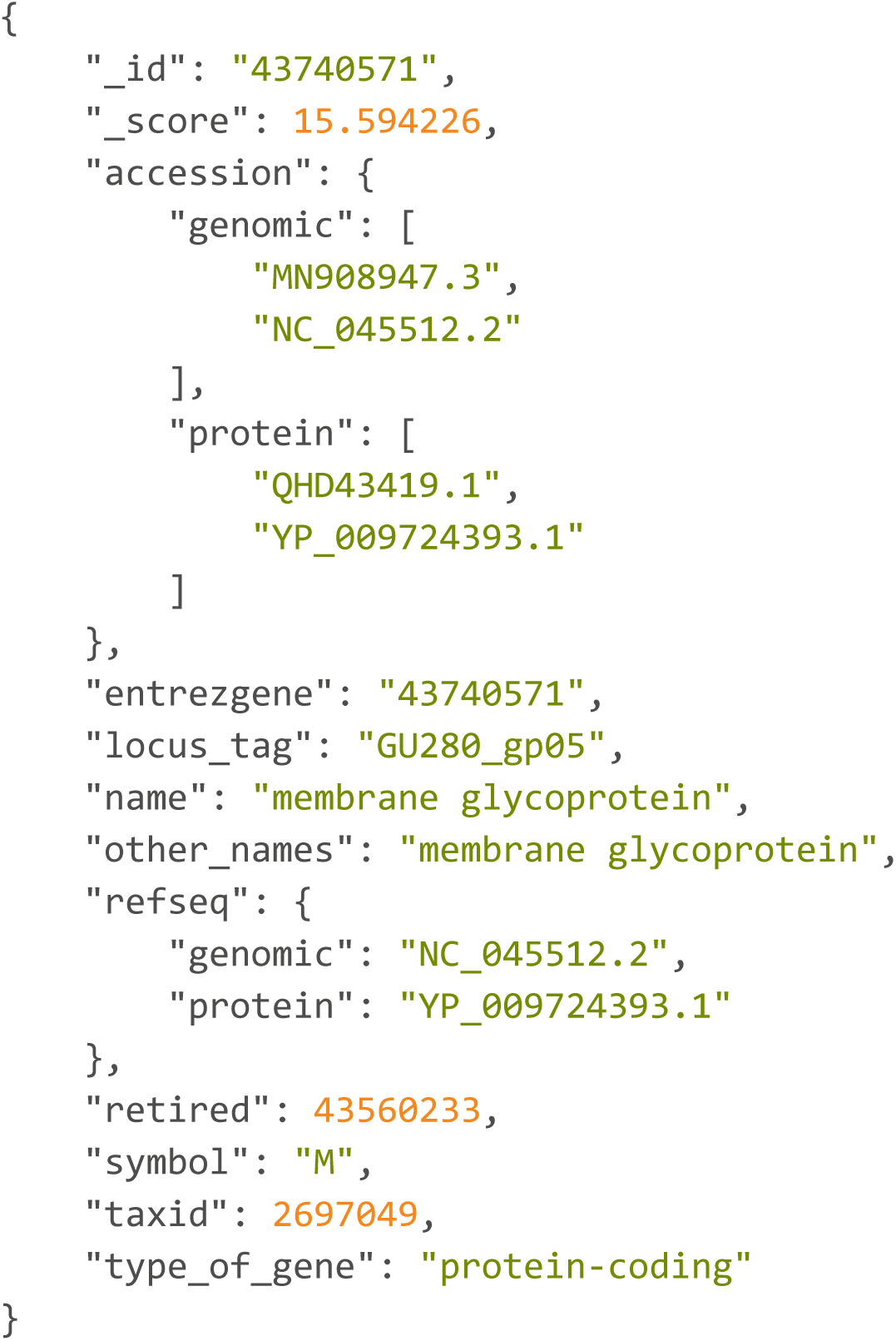
JavaScript Object notation output of the mygene,info output for gene with NCBI gene identifier 43740571.

### UniProt

The annotations retrieved from mygene.info also contain protein identifiers such as UniProt, RefSeq and PDB, however, their respective names are lacking. To obtain names and mappings to other protein identifiers, RefSeq and UniProt were consulted. Refseq annotations were acquired using the earlier mentioned NCBI EUtils. UniProt identifiers are acquired using the SPARQL endpoint of UniProt, which is a rich resource for protein annotations provided by the Swiss Bioinformatics Institute. Figure 5 shows the SPARQL query that was applied to acquire the protein annotations.

**Figure 5:**
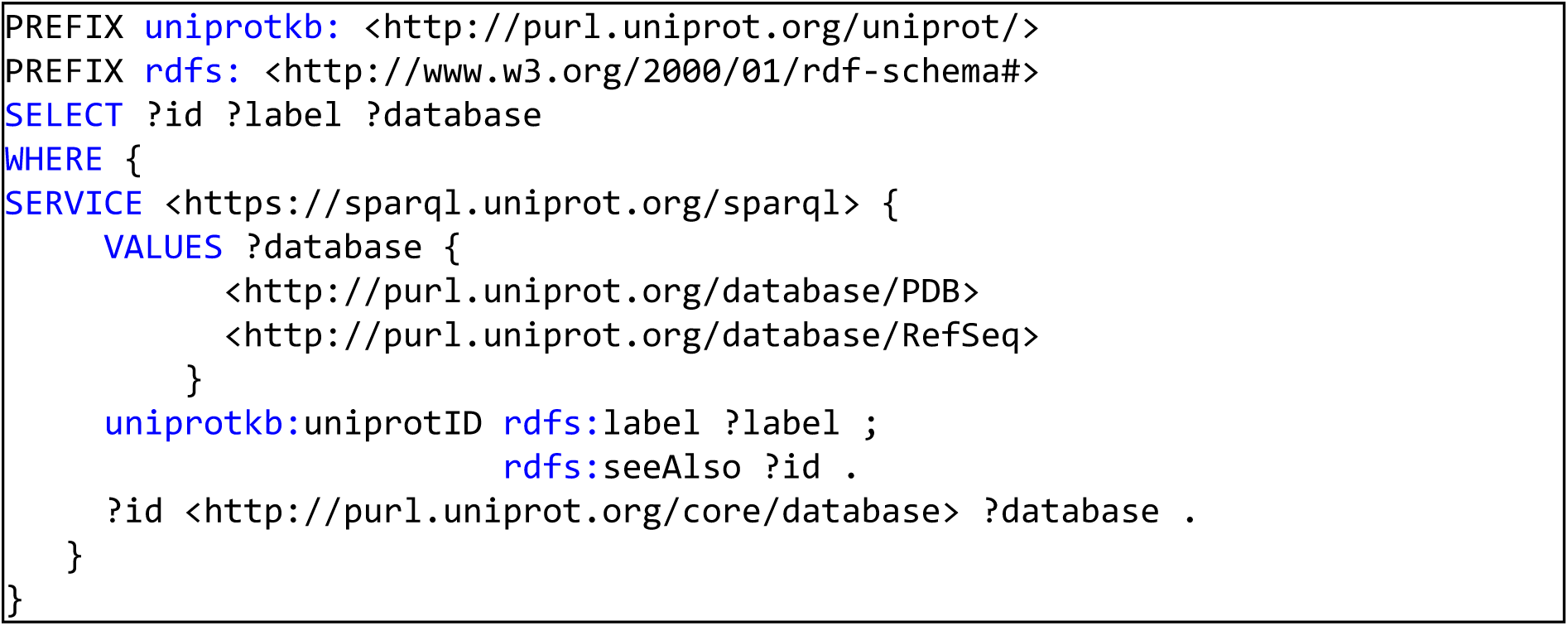
The UniProt SPARQL query used to obtain additional protein annotations, descriptions and external resources.

### Reconciliation with Wikidata

Before the aggregated information on viruses, genes and proteins can be added to Wikidata, reconciliation with Wikidata is necessary. If Wikidata items exist they are updated, otherwise, new items are created. Reconciliation is driven by mapping existing identifiers in both the primary resources and Wikidata. It is possible to reconcile based on strings, but this is dangerous due to the ambiguity of the labels used (37). When items on concepts are added to Wikidata that lack identifiers overlapping with the primary resource, reconciliation is challenging. Based on the Shape Expressions, the following properties are identified for use in reconciliation. For Proteins these are Uniprot ID (P352) and RefSeq protein ID (P637). For genes these are NCBI Gene ID (P351) and Ensembl Gene (ID) P594. When sourced information matches none of these properties, then a new item is created, if the concepts from the primary source reconcile with Wikidata items these are updated.

### Wikidataintegrator

Wikidata integrator is a Python library (https://github.com/SuLab/WikidataIntegrator) that wraps around the Wikidata API (https://www.wikidata.org/w/api.php) (10). From external resources such as the NCBI, gene and taxonomy statements have been compiled with provenance and assigned to the associated Wikidata items. When an item did not exist (or was not recognized), it was created. The module compiled a list of statements by parsing the primary sources under scrutiny and extracted what statements already existed on Wikidata. A JavaScript Object Notation (JSON) string was created that resembled the JSON data model used by the Wikidata API. This JSON string was then submitted to the Wikidata API for ingestion.

### Data Integration Use Cases / Validation

#### WikiPathways and BridgeDb

WikiPathways is a biological pathway database and can visualize the details of interactions between genes, proteins, metabolites, and other entities participating in biological processes. It depends on BridgeDb to map identifiers of external data and knowledge to the identifiers used for the genes, proteins, and metabolites in the pathways (38). Furthermore, mappings to Wikidata are required to establish the link of biological entities in pathways and journal articles that have those entities as main topics. Therefore, the virus genes and proteins are required to exist in Wikidata, enabling the interoperability between WikiPathways and Wikidata. Additionally, new virus mapping databases for BridgeDb are created by extracting the new virus gene and protein data, including links between Wikidata, NCBI Gene, RefSeq, UniProt and Guide to Pharmacology Target identifiers using a SPARQL query (https://github.com/bridgedb/Wikidata2Bridgedb). The mapping databases will be updated regularly and will allow pathway curators to annotate virus genes and proteins in their pathways and provide link outs on the WikiPathways website.

The COVID-19 related pathways from WikiPathways COVID-19 Portal are added to Wikidata using the approach previously described (10). For this, a dedicated repository has been set up to hold the GPML files, the internal WikiPathways file format, The GPML is converted into RDF files with the WikiPathways RDF generator (39), while the files with author information are manually edited. For getting the most recent GPML files, a custom Bash script was developed (getPathways.sh in the SARS-CoV-2-WikiPathways repository). The conversion of the GPML to RDF uses the previously published tools for WikiPathways RDF (39). Here, we adapted the code with a unit test that takes the pathways identifier as parameter. This test is available in the SARS-CoV-2-WikiPathways branch of GPML2RDF along with a helper script (createTurtle.sh). Based on this earlier generated pathway RDF and using the Wikidataintegrator library, the WikiPathways bot was used to populate Wikidata with additional statements and items. The pathway bot was extended with the capability to link virus proteins to the corresponding pathways, which was essential to support the Wikidata resource. These changes can be found in the *sars-cov-2-wikipathways-2* branch.

#### Scholia

The second use case is to demonstrate how we can link virus gene and protein information to literature. Here, we used Scholia (https://scholia.toolforge.org/) as a central tool (13). It provides a graphical interface around data in Wikidata, for example, literature about a specific coronavirus protein (e.g. Q87917585 for the SARS-CoV-2 spike protein). Scholia uses SPARQL queries to provide information about topics. We annotated literature around the HCoVs with the specific virus strains, the virus genes, and the virus proteins as ‘main topic’.

## Results

### Semantic data landscape

To align the different sources in Wikidata, a common data schema is needed. We have created a collection of schemas that represent the structure of the items added to Wikidata. Input to the workflow is the NCBI taxon identifier, which is input to mygene.info (see Figure 3). Taxon information is obtained and added to Wikidata according to a set of linked Entity Schemas (<u>E170</u>: virus, <u>E174</u>: strain, <u>E69</u>: disease). Gene annotations are obtained and added to WIkidata following the Schemas (<u>E165</u>: virus gene, <u>E169</u>: virus protein) and protein annotations are obtained and added to Wikidata following the two schemas. The last two schemas are an extension from more generic schemas for proteins (<u>E167</u>) and genes (<u>E75</u>).

### Bots

The bots developed and used in this protocol are adaptations of the bots developed in the Gene Wiki project. On regular intervals, the bots run to update viral gene and protein annotations as well as pathway updates from WikiPathways. For the gene and protein annotations, we have also made a Jupyter Notebook. The bot that synchronizes virus, gene and protein information and the Jupyter Notebook are available at https://github.com/SuLab/Gene_Wiki_SARS-CoV. The bot that synchronizes the WikiPathways pathways with Wikidata was updated from the original version to allow adding proteins annotated with Wikidata identifiers and no longer requires pathways to be part of the WikiPathways *Curated Collection*. The customized bot source code is available at https://github.com/SuLab/scheduled-bots/blob/SARS-CoV-2/scheduled_bots/wikipathways/bot.py.

Both bots are now part of the automation server used in the Gene Wiki project. This runs on the Jenkins platform (40). Although Jenkins is mainly aimed at software deployments, its extended scheduling capabilities the synchronization procedure at set intervals that can be changed depending on the update speed of the external resources. The Jenkins jobs are available from http://jenkins.sulab.org/.

### Data added

Using the gene and proteins bots explained in the Methods section, missing genes and proteins have been added for the seven human coronaviruses. The results are summarized in Table 1. The automatically added and updated gene and protein items were manually curated. For SARS-CoV-2 all items were already manually created, and the bot only edited gene items. Thirteen out of 27 protein entries were created by the authors and saw edits, up to May 23, by up to five other Wikidata editors including two bots (Krbot and Edoderoobot). For the other species, all gene entries and most protein entries have been created by the bot. Only for MERS and SARS-CoV-2 some protein entries were added manually, including some by us. During this effort, which took three weeks, the bot created a number of duplicates. These have been manually corrected. It should also be noted that for SARS-CoV-2 many proteins and protein fragments do not have RefSeq or UniProt identifiers, mostly for the virus protein fragments.

**Table 1:** Summary of the seven human coronaviruses, including taxon identifiers, the Wikidata items, and the number of genes and proteins. The latter two are generated by the SPARQL queries `geneCount.rq` and `proteinCount.rq` in the supporting information.

| Virus strain | NCBI Taxon ID | Wikidata Qid | # Genes | # Proteins |
| --- | --- | --- | --- | --- |
| SARS-virus | 694009 | Q278567 | 14 | 11 |
| Middle East respiratory syndrome coronavirus | 1335626 | Q4902157 | 11 | 9 |
| Human Coronavirus NL63 | 277944 | Q8351095 | 7 | 6 |
| Human coronavirus 229E | 11137 | Q16983356 | 8 | 8 |
| Human coronavirus HKU1 | 290028 | Q16983360 | 9 | 9 |
| Human coronavirus OC43 | 31631 | Q16991954 | 9 | 8 |
| SARS-CoV-2 | 2697049 | Q82069695 | 11 | 27 |

### Use Cases

#### BridgeDb

Using the dedicated code to create a BridgeDb identifier mapping database for coronaviruses, mappings were extracted from Wikidata with a SPARQL query for the seven human coronaviruses and the SARS-related viruses. This resulted in a mapping database with 554 mappings between 368 identifiers (version 2020-05-27). This includes 170 Wikidata identifiers, 70 NCBI Gene identifiers, 60 UniProt identifiers, 58 RefSeq identifiers and 10 Guide to Pharmacology Target identifiers. The mapping file has been released on the BridgeDb website (https://bridgedb.github.io/data/gene_database/) and archived on Zenodo (41). The mapping database has also been loaded on the BridgeDb webservice at http://webservice.bridgedb.org/ which means it can be used in the next use case: providing links out for WikiPathways.

#### WikiPathways

The WikiPathways project is involved in an international collaboration to curate knowledge about the biological processes around SARS-CoV-2 and COVID-19. The authors have started a pathway specifically about SARS-CoV-2 (wikipathways:WP4846). To ensure interoperability, WikiPathways uses BridgeDb and taking advantage of the enriched BridgeDb webservice, WikiPathways now links out for HCoV genes and proteins (depending on availability of mappings) to RefSef, NCBI Gene, UniProt, and Scholia (see Figure 7). The latter links to the next use case, and provides a link to literature about the virus. It should be noted that for each gene and protein two Wikidata identifiers with links may be given. In that case, one is for the gene and one for the protein.

**Figure 6:**
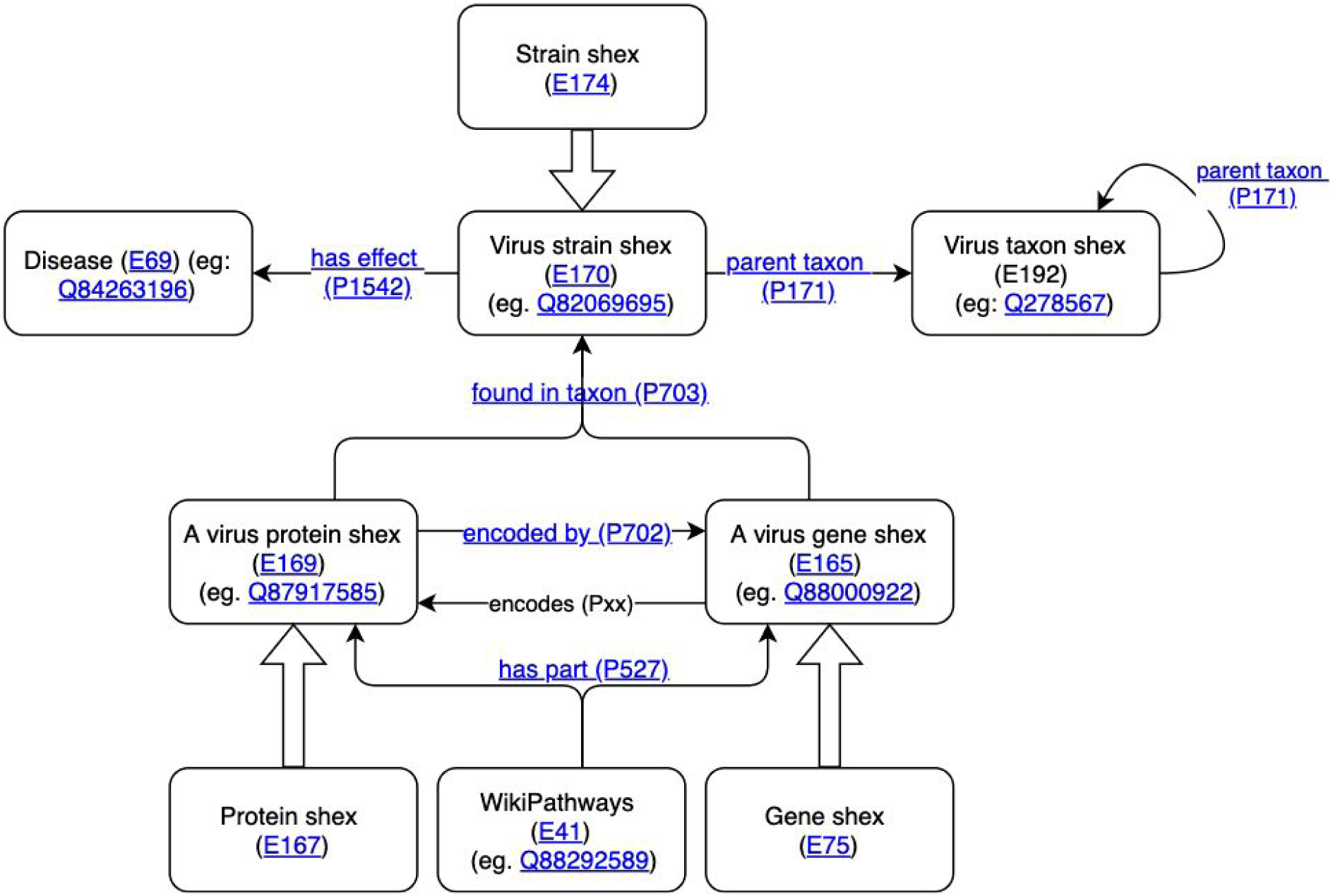
Overview of the ShEx schemas and the relations between them. All shapes, properties and items are available from within Wikidata.

**Figure 7:**
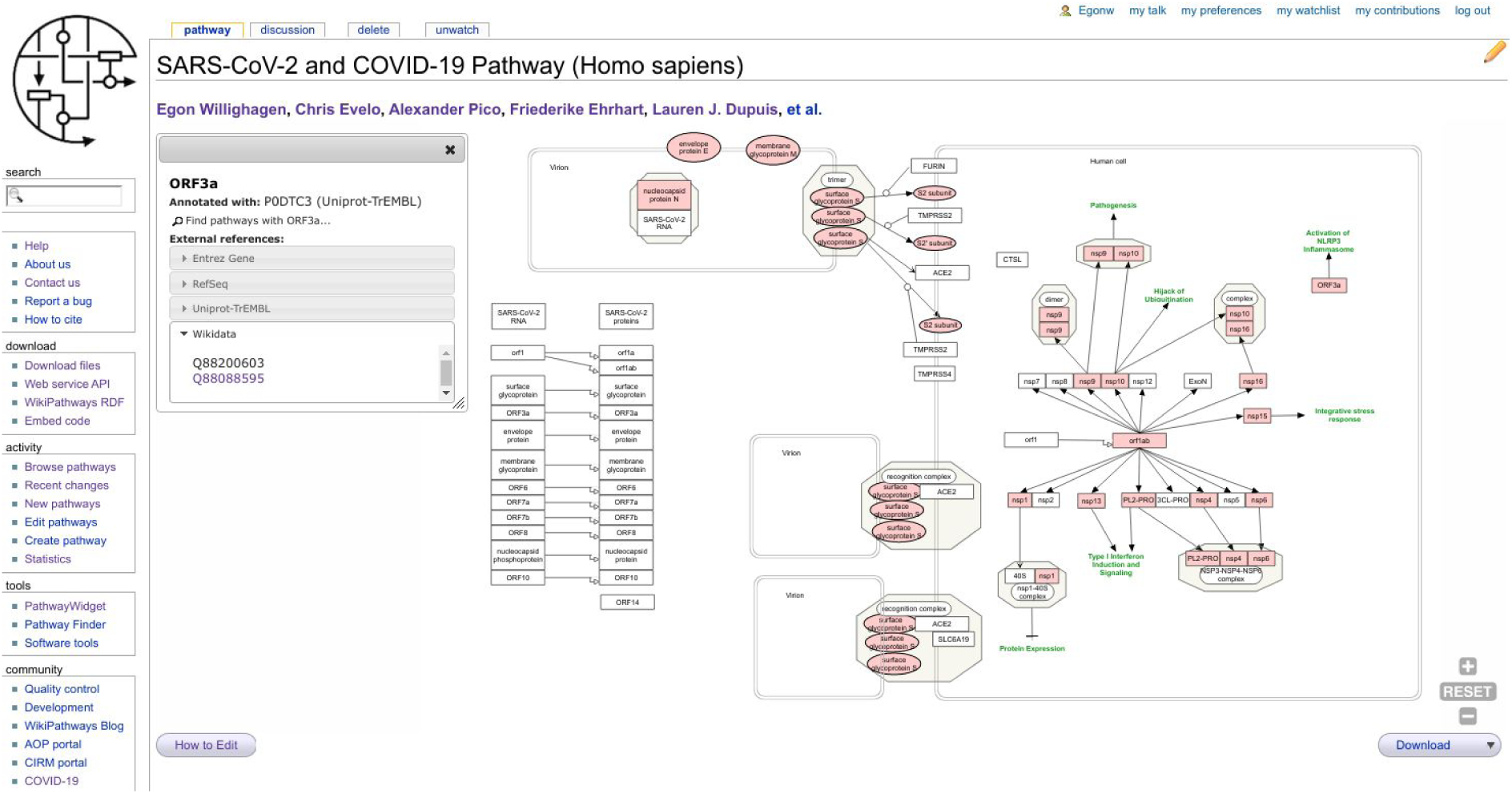
Screenshot of SARS-CoV-2 and COVID-19 Pathway in WikiPathways (wikipathways:WP4846) showing the BridgeDb popup box for the ORF3a protein, showing a link out to Scholia via the protein and gene’s Wikidata identifiers.

#### Scholia

The WikiPathways use case shows us that literature describes our knowledge about how coronaviruses work at a rather detailed level. Indeed, many articles discuss the genetics, homology of genes and proteins across viruses, or the molecular aspects of how these proteins are created and how they interfere with the biology of the human cell. The biological pathways show these processes, but ultimately the knowledge comes from literature. Wikidata allows us to link literature to specific virus proteins and genes, depending on what the article describes. For this it uses the ‘main subject’ property (<u>P921</u>). We manually annotated literature with the Wikidata items for specific proteins and genes. We developed two SPARQL queries to count the number of links between genes (https://w.wiki/Lsp) and proteins (https://w.wiki/Lsq) and the articles that discuss them. Scholia takes advantage of the ‘main subject’ annotation, allowing the creation of “topic” pages for each protein. For example, Figure 8 shows the topic page of the SARS-CoV-2 spike protein.

**Figure 8:**
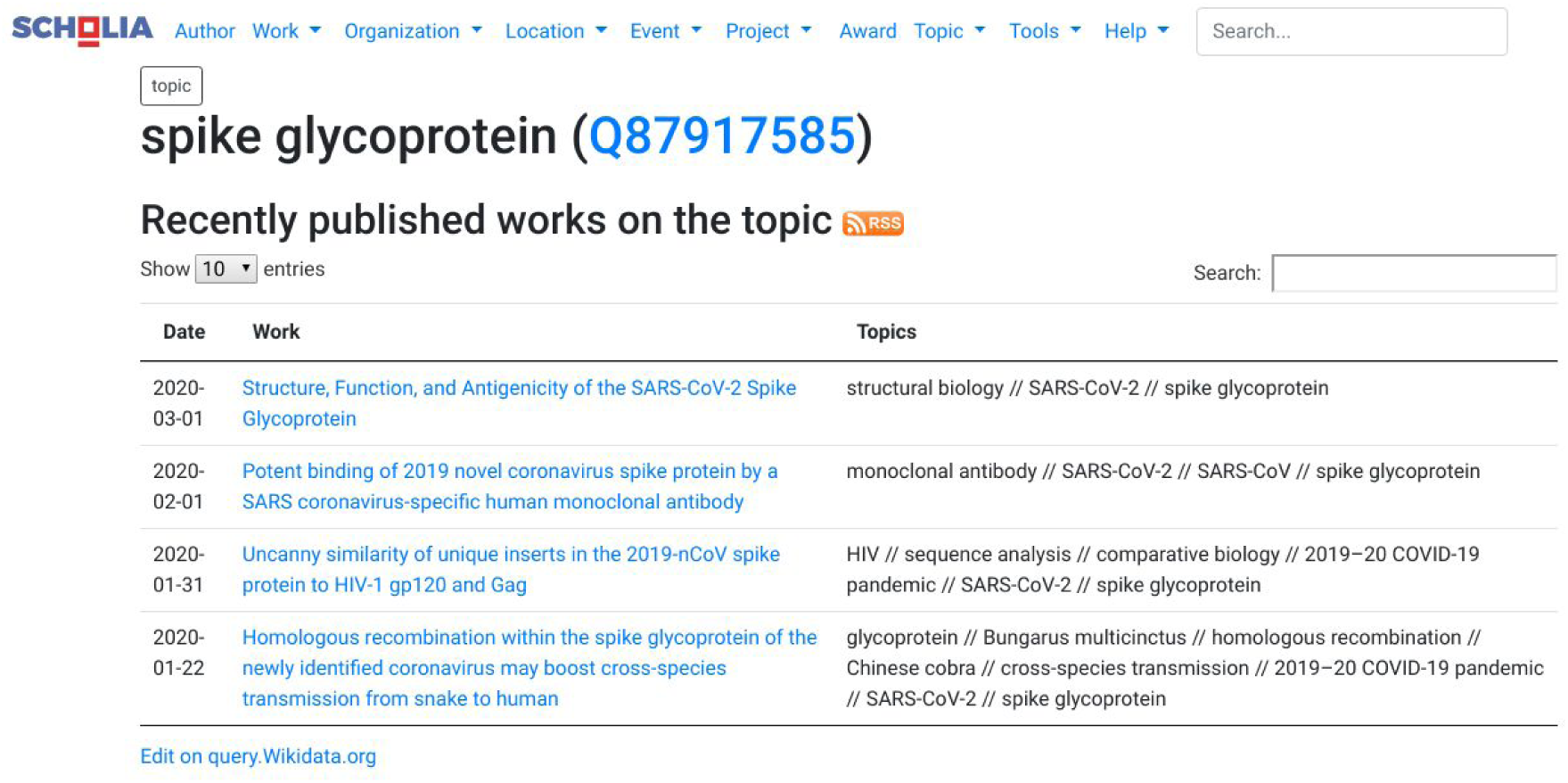
Screenshot of the Scholia page for the SARS-CoV-2 spike glycoprotein, it shows four articles that specifically discuss this protein.

## Discussion

This paper describes a protocol we developed to align genetic annotations from reference resources with Wikidata. Numerous annotations are scattered across different sources without any overall integration, thereby reducing the reusability of knowledge from different sources. Integration of the annotations from these resources is a complex and time consuming task. Each resource uses different ways to access the data from a user and machine perspective. Making use of these protocols programmatically to access and retrieve data of interest, requires the knowledge of various technologies and procedures to extract the information of interest.

Wikidata provides a solution. It is part of the semantic web, taking advantage of its reification of the Wikidata items as RDF. Data in Wikidata itself is frequently, often almost instantaneously, synchronized with the RDF resource and available through its SPARQL endpoint (http://query.wikidata.org). The modelling process turns out to be an important aspect of this protocol. Wikidata contains numerous entity classes as entities and more than 7000 properties which are ready for (re-)use. However, that also means that this is a confusing landscape to navigate. The ShEx Schema has helped us develop a clear model. This is a social contract between the authors of this paper, as well as documentation for future users.

Using these schemas, it was simpler to validate the correctness of the updated bots to enter data in Wikidata. The bots have been transferred to the Gene Wiki Jenkins platform. This allows the bots to be kept running regularly, pending on the ongoing efforts of the coronavirus and COVID-19 research communities. While the work of the bots will continue to need human oversight, potentially to correct errors, it provides a level of scalability and generally alleviates the authors from a lot of repetitive work.

One of the risks of using bots, is the possible generation of duplicate items. Though this is also a risk in manual addition of items, humans can apply a wider range of academic knowledge to resolve these issues. Indeed, in running the bots, duplicate Wikidata items were created, for which an example is shown in Figure 9. The Wikidataintegrator library does have functionality to prevent the creation of duplicates by comparing properties, based on used database identifiers. However, if two items have been created using different identifiers, these cannot be easily identified.

**Figure 9:**
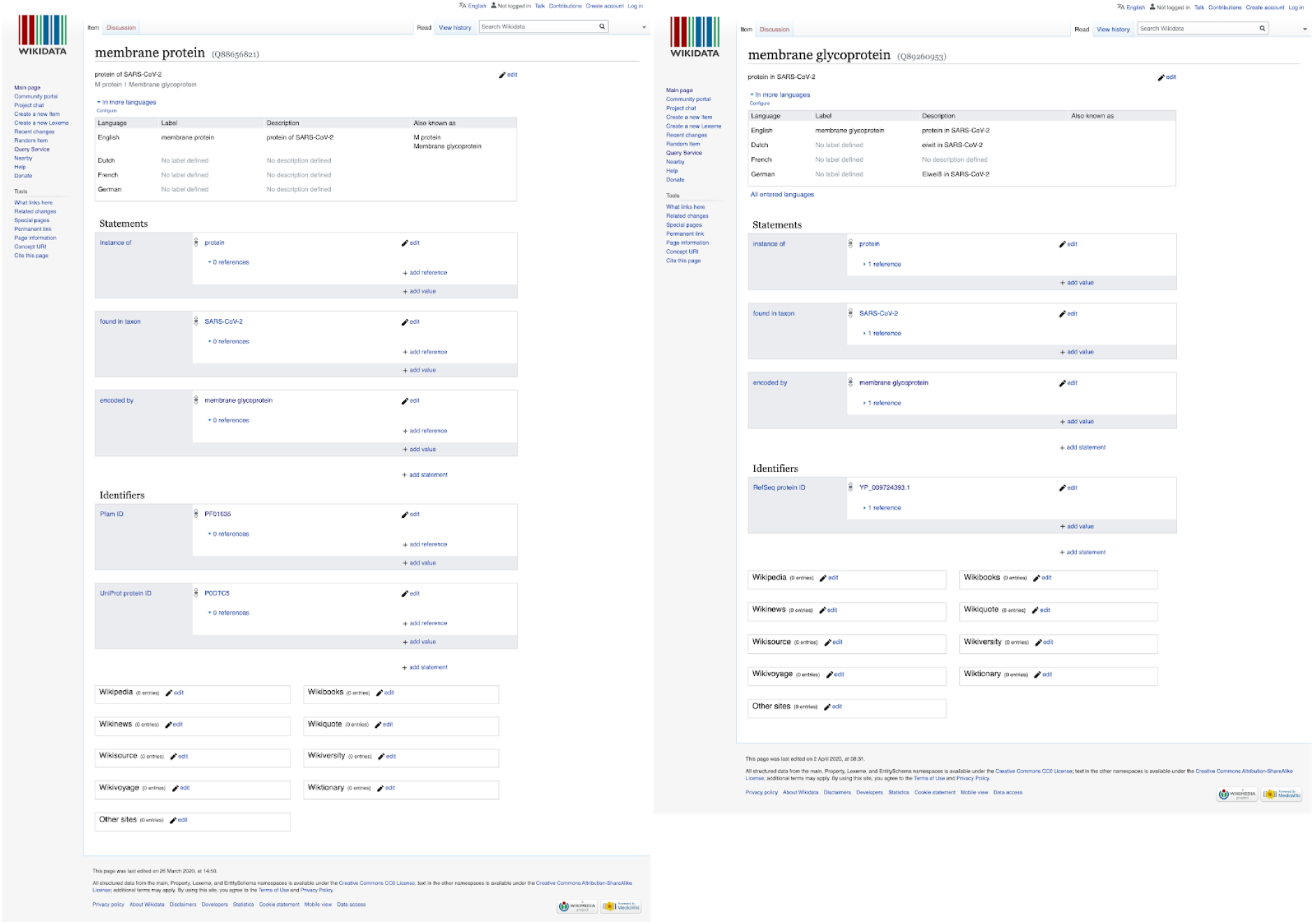
Comparison of two Wikidata entries for the SARS-CoV-2 membrane protein. An overlap between a Wikidata item and a concept from a primary source needs to have some overlap to allow automatic reconciliation. If there is no overlap, duplicates will be created and left for human inspection. Since this screenshot was made, the entries have been merged in a manually curation process.

Close inspection of examples, such as the one in Figure 9, showed that the duplicates were created because there was a lack of overlap between the data to be added and the existing item. The UniProt identifier did not yet resolve, because it was manually extracted from information in the March 27 pre-release (but now part of the regular releases). In this example, the Pfam protein families database (42) identifier was the only identifier upon which reconciliation could happen. However, that identifier pointed to a webpage that did not contain mappings to other identifiers. In addition, the lack of references to the primary source hampers the curator’s ability to merge duplicate items and expert knowledge was essential to identify the duplication. Fortunately, names used for these RNA viruses only refer to one protein as the membrane protein. Generally, the curator would have to revert to the primary literature to identify the overlap. Statements about ‘encoded by’ to the protein coding genes were found to be helpful as well. Reconciliation might be possible through sequence alignment, which means substantial expert knowledge and skills are required.

This makes reconciliation in Wikidata based on matching labels, descriptions and synonyms, matching statements and captured provenance (qualifiers and references) hazardous, due to different meanings to the same label. A targeted curation query (*geneAndProteinCurationQuery*.*rq*, see Supporting Information) was developed to highlight such duplications and manually curated seven duplicate protein entries for SARS-CoV-2 alone. This duplication is common and to be expected, particularly in rapidly evolving situations like a pandemic, when many groups contribute to the same effort. In this case, this paper only represents one group contributing to the *Wikidata:WikiProject COVID-19 (https://www.wikidata.org/wiki/Wikidata:WikiProject_COVID-19*). We also discovered that virus taxonomy is different from zoological taxonomy. For example, there is no clear NCBI taxon identifier for SARS-CoV-1 and after consultation with other projects, we defaulted to using the taxon identifier for the SARS-related CoVs, something that NCBI and UniProt seem to have done as well. Finally, we note that during the two weeks this effort took place, several other resources introduced overviews, including dedicated COVID-19 portals from UniProt (https://covid-19.uniprot.org/) and the Protein DataBank in Europe (https://PDBe.org/covid19/).

## Conclusion

This manuscript presents a protocol to link information from disparate resources, including NCBI Taxonomy, NCBI Gene, UniProt, PubMed, and WikiPathways. Using the existing Wikidata infrastructure, we developed semantic schemas for virus strains, genes, and proteins; bots written in Python to add knowledge on genes and proteins of the seven human coronaviruses and linked them to biological pathways in WikiPathways and to primary literature, visualized in Scholia. We were able to do so in the period of two weeks, using an ad-hoc team from existing collaborations, taking advantage of the open nature of the components involved.

## Funding

**National Institute of General Medical Sciences (R01 GM089820)**

- Andra Waagmeester, Andrew I. Su

**Alfred P. Sloan Foundation (grant number G-2019-11458)**

- Egon Willighagen

**Spanish Ministry of Economy and Competitiveness (Society challenges: TIN2017-88877-R)**

- Jose Emilio Labra Gayo, Daniel Fernández Álvarez

**Netherlands Organisation for Scientific Research funded UNLOCK project (NRGWI.obrug.2018.005)**

- Jasper Koehorst, Peter Schaap

**SYNTHESYS+ a Research and Innovation action funded under H2020-EU.1.4.1.2. Grant agreement ID: 823827**.

- Quentin Groom

## Acknowledgments

The protocol presented in this work would not be as powerful if it was not for the works of many, which ensure that the fruits of our work are immediately interoperable with lots of other efforts, even though that integration was the whole purpose of developing this protocol. We thank the work of the larger Wikidata:WikiProject COVID-19 community in which we operated, Wikidata volunteers in general, and in particular Tiago Lubiana, Ralf Stephan, Daniel Mietchen, and Finn Nielsen. Second, the work is partly driven by a collaboration in the WikiPathways COVID-19 project and we particularly thank Anders Riutta for setting up a commit hook to the repository with WikiPathways COVID-19 pathways and the authors of these pathways.

## Author contributions

AW, EW, JK, conceptual design, code development and wrote the manuscript. MK, wrote the manuscript. JL, Software, Visualization, Writing - review and editing. DF, minor review, code support with shexer. QG: virus-taxonomy review, Writing - review and editing. PS, LM: review and editing. All authors read and approved the final manuscript.

## Supporting Information

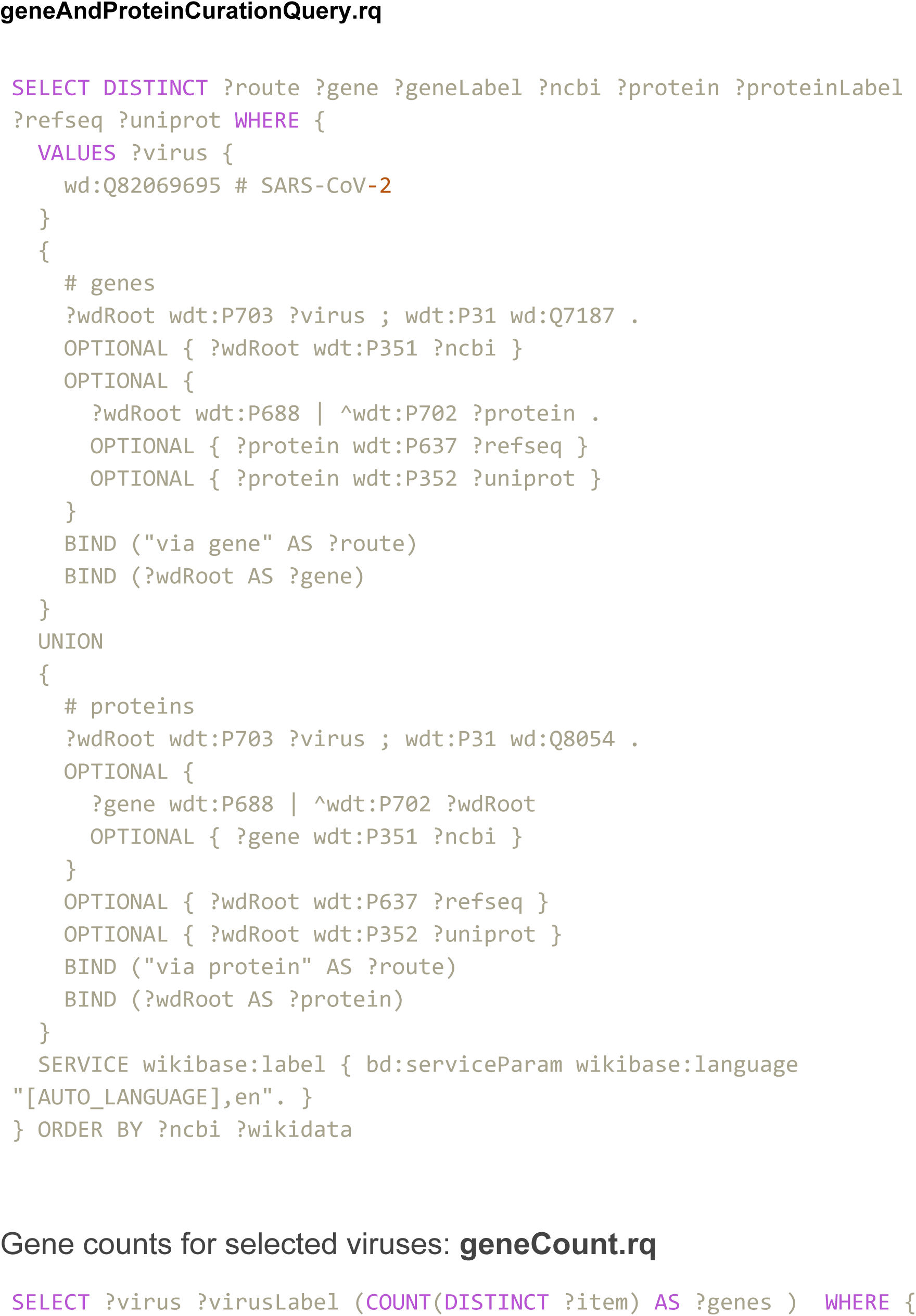
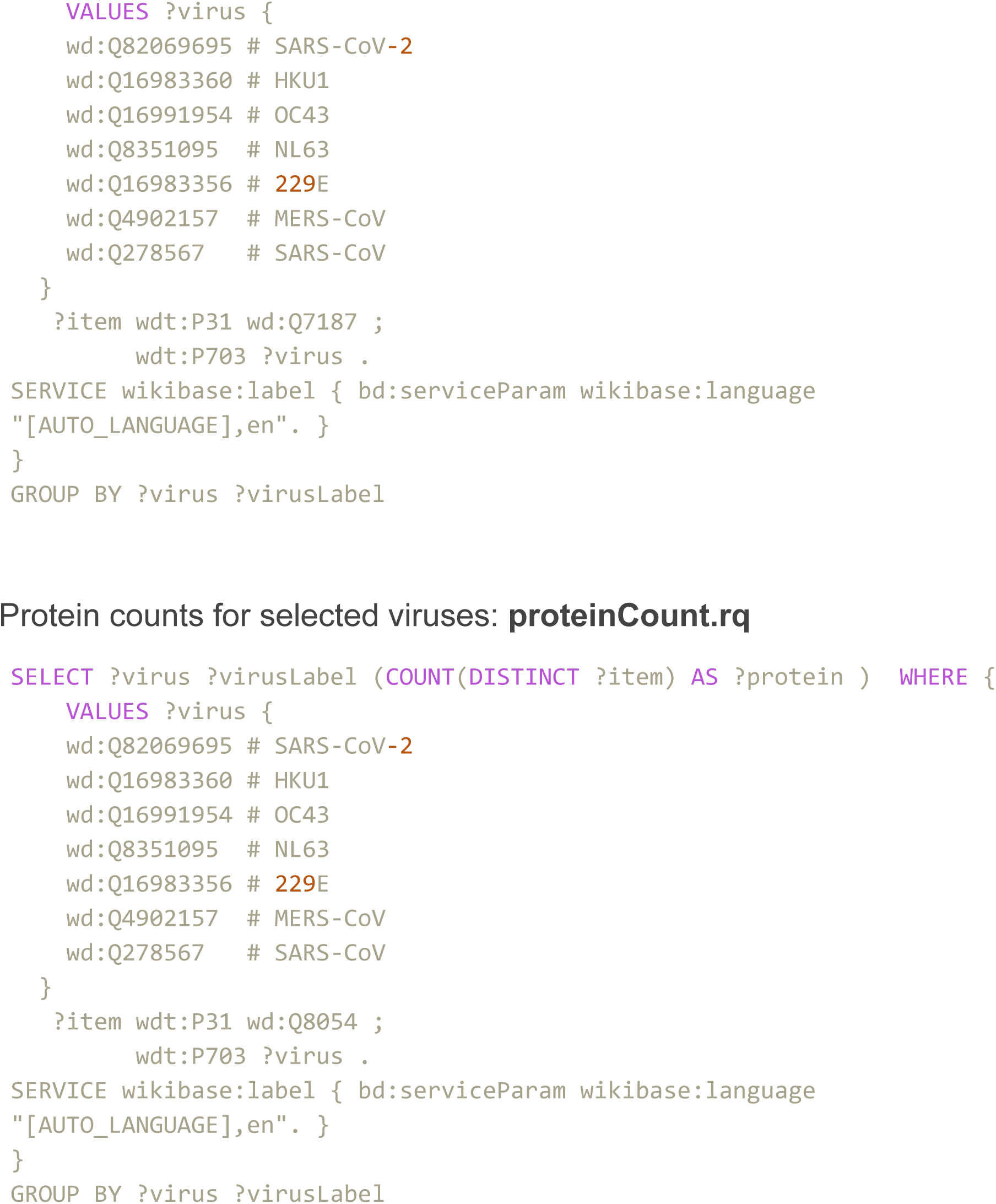

## References

1. Watkins J. Preventing a covid-19 pandemic. BMJ. 2020 Feb 28;m810.

2. outbreak.info [Internet]. Available from: https://github.com/SuLab/outbreak.info

3. Virus Outbreak Data Network (VODAN) [Internet]. Available from: https://www.go-fair.org/implementation-networks/overview/vodan/

4. CORD-19-on-FHIR -- Semantics for COVID-19 Discovery [Internet]. Available from: https://github.com/fhircat/CORD-19-on-FHIR

5. Ostaszewski M, Mazein A, Gillespie ME, Kuperstein I, Niarakis A, Hermjakob H, et al. COVID-19 Disease Map, building a computational repository of SARS-CoV-2 virus-host interaction mechanisms. Sci Data. 2020 Dec;7(1):136.

6. The Coronavirus and Open Science: Our reads and Open use cases [Internet]. 2020. Available from: https://sparceurope.org/coronaopensciencereadsandusecases/

7. Sondervan J, Bosman J, Kramer B, Brinkman L, Imming M, Versteeg A. The COVID-19 pandemic stresses the societal importance of open science. ScienceGuide [Internet]. 2020 Apr 3; Available from: https://www.scienceguide.nl/2020/04/dire-times-of-covid-19-stress-the-societal-importance-of-open-science/

8. Sharma M, Scarr S, Kelland K. Speed Science: The risks of swiftly spreading coronavirus research [Internet]. 2020 Feb. Available from: https://graphics.reuters.com/CHINA-HEALTH-RESEARCH/0100B5ES3MG/index.html

9. Mahase E. Covid-19: six million doses of hydroxychloroquine donated to US despite lack of evidence. BMJ. 2020 Mar 23;m1166.

10. Waagmeester A, Stupp G, Burgstaller-Muehlbacher S, Good BM, Griffith M, Griffith OL, et al. Wikidata as a knowledge graph for the life sciences. eLife. 2020 Mar 17;9:e52614.

11. Vrandečić D, Krötzsch M. Wikidata. Commun ACM. 2014 Sep;57(10):78–85.

12. Burgstaller-Muehlbacher S, Waagmeester A, Mitraka E, Turner J, Putman T, Leong J, et al. Wikidata as a semantic framework for the Gene Wiki initiative. Database. 2016 Mar;2016:baw015.#x002B;.

13. Nielsen FÅ, Mietchen D, Willighagen E. Scholia And Scientometrics With Wikidata. 2017 Sep 8 [cited 2018 Jun 26]; Available from: https://zenodo.org/record/1036595

14. Erxleben F, Günther M, Krötzsch M, Mendez J, Vrandečić D. Introducing Wikidata to the Linked Data Web. In: Mika P, Tudorache T, Bernstein A, Welty C, Knoblock C, Vrandečić D, et al., editors. The Semantic Web – ISWC 2014 [Internet]. Springer International Publishing; 2014. p. 50–65. (Lecture Notes in Computer Science; vol. 8796). Available from: http://dx.doi.org/10.1007/978-3-319-11964-9_4

15. Cyganiak R, Lanthaler M, Wood D. RDF 1.1 Concepts and Abstract Syntax. W3C; 2014 Feb.

16. Malyshev S, Krötzsch M, González L, Gonsior J, Bielefeldt A. Getting the Most Out of Wikidata: Semantic Technology Usage in Wikipedia’s Knowledge Graph. In: Vrandečić D, Bontcheva K, Suárez-Figueroa MC, Presutti V, Celino I, Sabou M, et al., editors. The Semantic Web – ISWC 2018 [Internet]. Cham: Springer International Publishing; 2018 [cited 2020 Apr 4]. p. 376–94. (Lecture Notes in Computer Science; vol. 11137). Available from: http://link.springer.com/10.1007/978-3-030-00668-6_23

17. Federhen S. The NCBI Taxonomy database. Nucleic Acids Res. 2012 Jan 1;40(D1):D136–43.

18. Brown GR, Hem V, Katz KS, Ovetsky M, Wallin C, Ermolaeva O, et al. Gene: a gene-centered information resource at NCBI. Nucleic Acids Res. 2015 Jan 28;43(D1):D36–42.

19. Consortium TU. UniProt: the universal protein knowledgebase. Nucleic Acids Res. 2017 Jan;45(D1):D158–D169.

20. wwPDB consortium, Burley SK, Berman HM, Bhikadiya C, Bi C, Chen L, et al. Protein Data Bank: the single global archive for 3D macromolecular structure data. Nucleic Acids Res. 2019 Jan 8;47(D1):D520–8.

21. Slenter DN, Kutmon M, Hanspers K, Riutta A, Windsor J, Nunes N, et al. WikiPathways: a multifaceted pathway database bridging metabolomics to other omics research. Nucleic Acids Res. 2018 Jan 4;46(D1):D661–D667.

22. Sayers EW, Beck J, Brister JR, Bolton EE, Canese K, Comeau DC, et al. Database resources of the National Center for Biotechnology Information. Nucleic Acids Res. 2020 Jan 8;48(D1):D9–16.

23. Thornton K, Solbrig H, Stupp GS, Labra Gayo JE, Mietchen D, Prud’hommeaux E, et al. Using Shape Expressions (ShEx) to Share RDF Data Models and to Guide Curation with Rigorous Validation. In: Hitzler P, Fernández M, Janowicz K, Zaveri A, Gray AJG, Lopez V, et al., editors. The Semantic Web [Internet]. Cham: Springer International Publishing; 2019 [cited 2020 Apr 4]. p. 606–20. (Lecture Notes in Computer Science; vol. 11503). Available from: http://link.springer.com/10.1007/978-3-030-21348-0_39

24. Prud’hommeaux E, Labra Gayo JE, Solbrig H. Shape expressions: an RDF validation and transformation language. In: Proceedings of the 10th International Conference on Semantic Systems - SEM ‘14 [Internet]. Leipzig, Germany: ACM Press; 2014 [cited 2020 Apr 5]. p. 32–40. Available from: http://dl.acm.org/citation.cfm?doid=2660517.2660523

25. Zhu N, Zhang D, Wang W, Li X, Yang B, Song J, et al. A Novel Coronavirus from Patients with Pneumonia in China, 2019. N Engl J Med. 2020 Feb 20;382(8):727–33.

26. Berners-Lee T, Hendler J, Lassila O. The Semantic Web. Sci Am. 2001 May;284(5):34–43.

27. Seaborne A, Harris S. SPARQL 1.1 Query Language. W3C; 2013 Mar.

28. W3C OWL Working Group. OWL 2 Web Ontology Language Document Overview [Internet]. 2012 Dec. Available from: https://www.w3.org/TR/owl2-overview/

29. Berners-Lee T. Linked Data [Internet]. 2006. Available from: https://www.w3.org/DesignIssues/LinkedData.html

30. Samwald M, Jentzsch A, Bouton C, Kallesøe C, Willighagen E, Hajagos J, et al. Linked open drug data for pharmaceutical research and development. J Cheminformatics. 2011;3(1):19.

31. Hernández D, Hogan A, Krötzsch M. Reifying RDF: What Works Well With Wikidata? In: Proceedings of the 11th International Workshop on Scalable Semantic Web Knowledge Base Systems co-located with 14th International Semantic Web Conference (ISWC 2015) [Internet]. Bethlehem, PA, USA; 2015. (CEUR Workshop Proceedings). Available from: http://ceur-ws.org/Vol-1457/SSWS2015_paper3.pdf

32. Jose Emilio Labra Gayo. RDFSHape: Online demo implementation of ShEx and SHACL [Internet]. Zenodo; 2018 [cited 2020 Apr 4]. Available from: https://zenodo.org/record/1412128

33. Pablo Menéndez Suárez, Jose Emilio Labra Labra Gayo. YaShE [Internet]. Zenodo; 2020 [cited 2020 Apr 4]. Available from: https://zenodo.org/record/3739108

34. Fernández-Álvarez D, García-González H, Frey J, Hellmann S, Gayo JEL. Inference of Latent Shape Expressions Associated to DBpedia Ontology. In: International Semantic Web Conference (P&D/Industry/BlueSky). 2018.

35. Sayers E. E-utilities Quick Start [Internet]. NCBI; 2018. Available from: https://www.ncbi.nlm.nih.gov/books/NBK25500/

36. Wu C, MacLeod I, Su AI. BioGPS and MyGene.info: organizing online, gene-centric information. Nucleic Acids Res. 2013 Jan 1;41(D1):D561–5.

37. Mungall C. Never mind the logix: taming the semantic anarchy of mappings in ontologies [Internet]. Monkeying around with OWL. 2019. Available from: https://douroucouli.wordpress.com/2019/05/27/never-mind-the-logix-taming-the-semantic-anarchy-of-mappings-in-ontologie/

38. van Iersel MP, Pico AR, Kelder T, Gao J, Ho I, Hanspers K, et al. The BridgeDb framework: standardized access to gene, protein and metabolite identifier mapping services. BMC Bioinformatics. 2010 Jan;11(1):5+.

39. Waagmeester A, Kutmon M, Riutta A, Miller R, Willighagen EL, Evelo CT, et al. Using the Semantic Web for Rapid Integration of WikiPathways with Other Biological Online Data Resources. Ouzounis CA, editor. PLOS Comput Biol. 2016 Jun 23;12(6):e1004989.

40. Armenise V. Continuous Delivery with Jenkins: Jenkins Solutions to Implement Continuous Delivery. In: 2015 IEEE/ACM 3rd International Workshop on Release Engineering [Internet]. Florence, Italy: IEEE; 2015 [cited 2020 May 31]. p. 24–7. Available from: http://ieeexplore.ieee.org/document/7169448/

41. Kutmon, Martina, Willighagen, Egon. BridgeDb: Human and SARS-related corona virus gene/protein mapping database derived from Wikidata [Internet]. Zenodo; 2020 [cited 2020 May 31]. Available from: https://zenodo.org/record/3860798

42. El-Gebali S, Mistry J, Bateman A, Eddy SR, Luciani A, Potter SC, et al. The Pfam protein families database in 2019. Nucleic Acids Res. 2019 Jan 8;47(D1):D427–32.

